# Single-nuclei RNA sequencing uncovers non-cell autonomous changes in cerebellar astrocytes and oligodendrocytes that may contribute to Spinocerebellar Ataxia Type 1 (SCA1) pathogenesis

**DOI:** 10.1101/2021.10.28.466301

**Authors:** Ella Borgenheimer, Ying Zhang, Marija Cvetanovic

## Abstract

Glial cells, including astrocytes and oligodendrocytes are important for normal brain function. In many neurodegenerative diseases glial cells undergo significant morphological, functional and gene expression changes termed reactive gliosis. The cause, identity and neuroprotective or neurotoxic nature of these changes remains incompletely understood. This knowledge in needed to develop a framework of how individual pathological changes in glial cells contribute to progressive dysfunction and selective neuronal vulnerability in neurodegenerative diseases. This is particularly relevant during the early disease stages that allow for the effective therapies and reversal or slowing of disease phenotypes. Spinocerebellar ataxia type 1 (SCA1) is a progressive neurodegenerative disease caused by an abnormal expansion of CAG repeats in the gene *Ataxin1* (*ATXN1*). While mutant ATXN1 is expressed broadly throughout the brain, SCA1 is characterized by severe degeneration of cerebellar Purkinje cells (PCs). Despite major advances in dissecting the effects of mutant ATXN1 on Purkinje cells, much less is understood how cerebellar astrocytes and oligodendrocytes respond to and contribute to Purkinje cell dysfunction in SCA1. To address this question we performed cerebellar single nuclei RNA sequencing (snRNA seq) of early disease stage *Pcp2-ATXN1[82Q]* mice, a transgenic SCA1 mouse model expressing mutant ATXN1 only in Purkinje cells. We found no changes in cell numbers in the SCA1 cerebellum. We validated previously indicated pathway and gene expression changes in the Purkinje cells, and identified novel DEG and pathways in Purkinje cells, including *Ralyl* that may provide compensatory roles and maintain PC function. Importantly we identified profound non-cell autonomous and potentially neuroprotective gene expression and pathway alterations in Bergman glia, velate astrocytes and oligodendrocytes that may contribute to disease pathogenesis.

## Introduction

Glia cells play key roles required for the normal brain function [1]. These roles include maintaining homeostasis of ions, neurotransmitters and water, providing neurotrophic and energy support, removal of unused synapses, and fast propagation of neuronal potentials [2]. In most if not all neurodegenerative diseases, glial cells undergo reactive gliosis, a process that includes molecular, morphological and functional changes [3][4]. These glial changes have been shown to play active role in the pathogenesis of neurodegenerative diseases [5][6]. In some situations, reactive gliosis can be beneficial and delay and ameliorate pathogenesis, but reactive glia have been also found to exacerbate pathogenesis [7][8][9][10]. The neuroprotective or toxic nature of reactive glial phenotypes likely depends on the stage of disease progression and the mechanism and intensity of signaling that leads to their activation.

Glia can be activated in response to distress signaling from neurons or other neural cells. In many neurodegenerative diseases a disease-causing genes and proteins are expressed in glial cell. Therefore, in these diseases glia can be also activated in response to the toxic effect of disease causing genes and proteins within the glial cells themselves. It is important to understand both glial cell –autonomous and non-cell-autonomous reactive responses, as well as how they interact to produce resulting glial phenotypes in disease. We propose that the way to start to address these questions is to reduce complexity and use disease models of neurodegeneration that express mutant proteins only in neurons to first understand the non-cell autonomous glial changes (that are in response to neuronal/neural distress and dysfunction).

Spinocerebellar ataxia type-1 (SCA1) is dominantly inherited neurodegenerative disorder caused by the abnormal expansion of CAG repeats in the *Ataxin-1* (*ATXN1)* gene [11]. Repeat expansions of 39 or more CAG repeats result in SCA1 disease with larger expansions leading to earlier onset and more severe symptoms [12]. Symptoms of SCA1 include loss of balance and coordination, swallowing and speech difficulties, impairments in cognition and mood, and premature death [13][14][15][16]. Selective neuronal vulnerability in different brain regions despite wide expression of mutant proteins is a feature shared by many neurodegenerative diseases. In case of SCA1 cerebellar Purkinje cells (PCs) are most affected although mutant ATXN1 is expressed throughout the brain. While *Atxn1*-targeting antisense oligonucleotides (ASO) show promise in pre-clinical trials [17], there are currently no disease-modifying treatments available for SCA1 [18], underlying the need for increased understanding of pathogenesis and neuronal vulnerability in SCA1. Severe vulnerability of PCs in SCA1 is likely brought about by the combination of the toxic effects of mutant ATXN1 within PCs and the PCs microenvironment, including glial cells. In this respect it is important to note that glial cells in the cerebellum have a very distinct transcriptome compared to the glial cells in other brain regions [19][20]. Uniqueness of cerebellar glia gets even more pronounced during ageing. Yet while neurodegeneration associated glial gene expression changes have been studied intensively in the other brain regions, less is known about gene expression changes in cerebellar glia during neurodegeneration [21][4].

Moreover, there are several excellent SCA1 mouse models that exhibit Purkinje cell pathology and motor deficits. In transgenic *Pcp2-ATXN1[82Q]* line mutant ATXN1 is expressed only in Purkinje cells in the cerebellum. Thereby, this mouse model is well suited to investigate the non-cell autonomous glial changes in neurodegenerative disease (that are in response to neuronal/neural distress and dysfunction)[22].

We decided to focus our study on Bergmann glia, velate astrocytes and oligodendrocytes for the following reasons. Bergmann glia are a special type of astrocytes whose cell bodies reside in Purkinje cell layer surrounding the cell bodies of Purkinje cells. BG processes extend into the molecular layer alongside PCs dendrites enveloping PCs synapses where BG regulate concentration of ions and neurotransmitters [23]. We have previously shown that Bergmann glia (BG) undergo reactive gliosis in patients with SCA1 and in SCA1 mouse models [24][25][26]. Moreover, we have demonstrated that early in disease progression reactive astrogliosis has a beneficial role delaying onset and slowing the progression of SCA1 pathogenesis in the *Pcp2-ATXN1[82Q]* line [25]. Velate astrocytes are unique among brain astrocytes as they are significantly outnumbered by granule neurons with glia to neuron ratio of ~ 0.2 while in the rest of the brain it can be up to 10. Cells of oligodendrocytes lineage are important for cerebellar development and motor learning [27][28]. Previous studies have found significant changes in the cerebellar white matter in SCA1 patients and in *Pcp2-ATXN1[82Q]* mice, indicating that oligodendrocytes may be affected in SCA1 [29][30]. Moreover, recent study found a decrease in oligodendrocyte (OL) numbers in human post-mortem samples from patients with SCA1 and altered expression of number of oligodendrocyte precursor cells (OPC)/OL genes in bulk RNA set from SCA1 knock-in mice *Atxn1^154Q/2Q^* line.

The bulk RNA sequencing has been useful for determining cerebellum-wide changes in the gene expression in mouse models of SCA1. However, bulk RNA sequencing precludes the detection of transcriptional changes at the single-cell level and is also confounded by the possible change in the proportion of cell types. Therefore to investigate molecular non-cell autonomous gene and pathways changes in cerebellar glia we have used single-nuclei RNA sequencing.

## Materials and Methods

### Mice

The creation of the *Pcp2-ATXN1[82Q]* mice was previously described [31]. We have backcrossed these mice onto C57BL/6 background for 10 generations. As CAG repeats are unstable and tend to shrink in mice we periodically sequence CAG region to evaluate number of repeats [32]. At the time of experiments the average number of CAG repeats in our colony was 71. Animal experimentation was approved by Institutional Animal Care and Use Committee (IACUC) of University of Minnesota and was conducted in accordance with the National Institutes of Health’s (NIH) Principles of Laboratory Animal Care (86–23, revised 1985), and the American Physiological Society’s Guiding Principles in the Use of Animals [33].

### Nuclei isolation

Nuclei for RNA sequencing were isolated using detergent mechanical lysis protocol as previously described [34]. Briefly, frozen or fresh whole cerebellum was placed into 1.5mL tube with 500uL pre-chilled detergent lysis buffer [low sucrose buffer (LSB) (sucrose 0.32M, 10 mM HEPES, 5mM CaCl2, 3mM MgAc, 0.1mM EDTA, 1mM DTT) with 0.1% Triton-X] and tissue was homogenized using mechanical homogenizer. 40um strainer was placed over a pre-chilled 50mL tube and pre-wetted with 1ml LSB. 1mL of LSB was added to the tube containing the crude nuclei in the lysis buffer and mixed gently by pipetting (2-3 times). Crude nuclei prep was then passed over the 40uM strainer into the pre-chilled 50mL tube, washed with 1 ml LSB and centrifuged at 3200g for 10 minutes at 4C The pellet was resuspended in 3mL of LSB, gently swirling to remove the pellet from the wall to facilitate the resuspension and left on ice for 2 min. Suspension was transferred to an Oak Ridge tube and nuclei were homogenized in LSB for 15-30s, keeping the sample on ice. Using a serological pipette, 12.5mL of density sucrose buffer (sucrose 1M, 10 mM HEPES, 3mM MgAc, 1mM DTT) was layered underneath the LSB homogenate, taking care not to create a bubble that disrupts the density layer. Samples were then centrifuge at 3200g for 20 minutes at 4C and pelleted nuclei were resuspended in the resuspension solution (PBS, 0.4 mg/ml BSA, 0.2 U/μl RNAse inhibitor). Nuclei were filtered through a 40um pore-size strainer followed by a 30um and 20um pore-size strainers. Small sample of nuclei was pelleted and added to the slide with VectaShield with DAPI to verify single nuclei isolation under fluorescent microscope. The nuclear suspensions were processed by the Genomic Core at the University of Minnesota using 10X Chromium 3' GEX Capture to Library Preparation (Chromium Next GEM Single Cell 3◻ Reagent Kits v3.1 with Single Cell 3’ v3.1 Gel Beads and Chromium Next GEM Chip G).

### Sequencing and analysis

Library quality control was performed using MiSeq system to estimate average numbers of nuclei per donor mouse. Then all nuclei from 6 donors were multiplexed and sequenced on two independent runs on the Illumina NovaSeq platform, using 1 full lane of S4 chip each time. Sequencing depth ranges from ~41,000 to ~120,000 reads per nuclei. Raw, demultiplexed fastq files were analyzed by CellRanger (v5.0.1) using reference genome mm10 (refdata-gex-mm10-2020-A), with the option that allows alignment to un-spliced pre-mRNAs (--include-introns). The resulted gene count table per donor was analyzed with R (v4.1.0) package Seurat (v 4.0.4). If bi-modal distribution was observed for nFeature (number of detected genes) per nuclei for a donor, nuclei composing the lower range peak were excluded from downstream analysis. Cell type identification was accomplished with SingleR (v 1.6.1) using DropViz (PMID: 30096299) Cerebellum MetaCells reference.

For identified cell types of Bergmann Cells, Velate Astrocytes, Granule Neurons, Purkinje Neurons and Oligodendrocytes, edgeR (v 3.34.1) was used to test differential abundance between control and KI donors. For the same identified cell types, scran (v 1.20.1) was used to normalize the single cell expression matrix, followed by limma-trend (v 3.48.3) to test for differential gene expression. Sample batches were considered as an independent factor in the design matrix of the statistical test (design = ~ genotype + batch). P values were adjusted using Benjamini-Hockberg method. Differential gene expression was determined by an adjusted p values of 0.05. The lists of differentially expressed genes from different cell types were further analyzed for GO term enrichment using ClusterProfiler (v 4.0.5).

Visualization of the results was generated using various R packages, including Seurat, ClusterProfiler, enrichplot (v 1.12.2), and ggplot2 (v 3.3.5).

### Immunofluorescent (IF) staining

IF was performed on a minimum of six different floating 45-μm-thick brain slices from each mouse (six technical replicates per mouse per region or antibody of interest). Confocal images were acquired using a confocal microscope (Olympus FV1000) using a 20X oil objective. Z-stacks consisting of twenty non-overlapping 1-μm-thick slices were taken of each stained brain slice per brain region (i.e., six z-stacks per mouse, each taken from a different brain slice). The laser power and detector gain were standardized and fixed between mice within a surgery cohort, and all images for mice within a cohort were acquired in a single imaging session to allow for quantitative comparison.

We used primary antibodies against Myein Basic protein, and GFAP. To quantify relative intensity of staining we measured the average signal intensity in the region of interest and normalized it to that of the WT littermate controls.

### Reverse transcription and quantitative polymerase chain reaction (RT-qPCR)

Total RNA was extracted from dissected mouse cerebella, medulla, and hippocampus using TRIzol (Life Technologies), and RT-qPCR was performed as described previously [8]. We used IDT Primetime primers for the following genes: *Ralyl*, *Mog*, *Ptch1*, *Ptch2*, *Gli* and *ApoE*. Relative mRNA levels were calculated using 18S RNA as a control and wild-type mice as a reference using 2^−ΔΔ-Ct^ as previously described.

### Data availability

All the data from this study are available from the authors.

## Results

### Mutant ATXN1 expression in Purkinje cells does not lead to the significant change in proportions of cerebellar cells

This study aimed to address question how neuronal dysfunction affects glial cells in cerebellar neurodegeneration. To ensure that glial cells are affected solely in the non-cell autonomous manner we have used Purkinje cells specific transgenic mouse model of SCA1, *Pcp2-ATXN1[82Q]* mice [22]. In these mice mutant *ATXN1* with 82 CAG repeats is selectively expressed under *Purkinje cell protein 2 (Pcp2)* promoter in cerebellar Purkinje cells. Because repeat length in trinucleotide repeat expansions is unstable and prone to further expansion or shrinkage [32], we routinely perform genetic sequencing of our mouse lines. We found that the number of repeats has recently decreased in our colony from 82 CAG to 71 CAG. Despite this reduction in CAG number, expression of Purkinje cell genes associated with SCA1 degeneration [35] remained significantly downregulated (**Supplementary Figure 1**).

Previous studies have shown advantages of single nuclei over single cell RNAseq. These include preservation of a larger number of cells across multiple subtypes, while minimizing the effects of cell dissociation on gene expression, and enriching for transcripts that are being actively transcribed *in vivo* [36][37]. Therefore we have performed RNA sequencing on isolated cerebellar individual nuclei that were captured and barcoded (**Figure 1A**)[38][39]. We isolated nuclei from the cerebella of six 12 weeks-old SCA1 and littermate control mice. We chose 12 weeks of age to capture the early stage when disease is still reversible [40] and as such therapeutically noteworthy. Bulk RNA seq showed significant gene expression changes in the cerebella at 12 weeks, yet there was no detectable PCs death [35][41].

**Figure 1.**
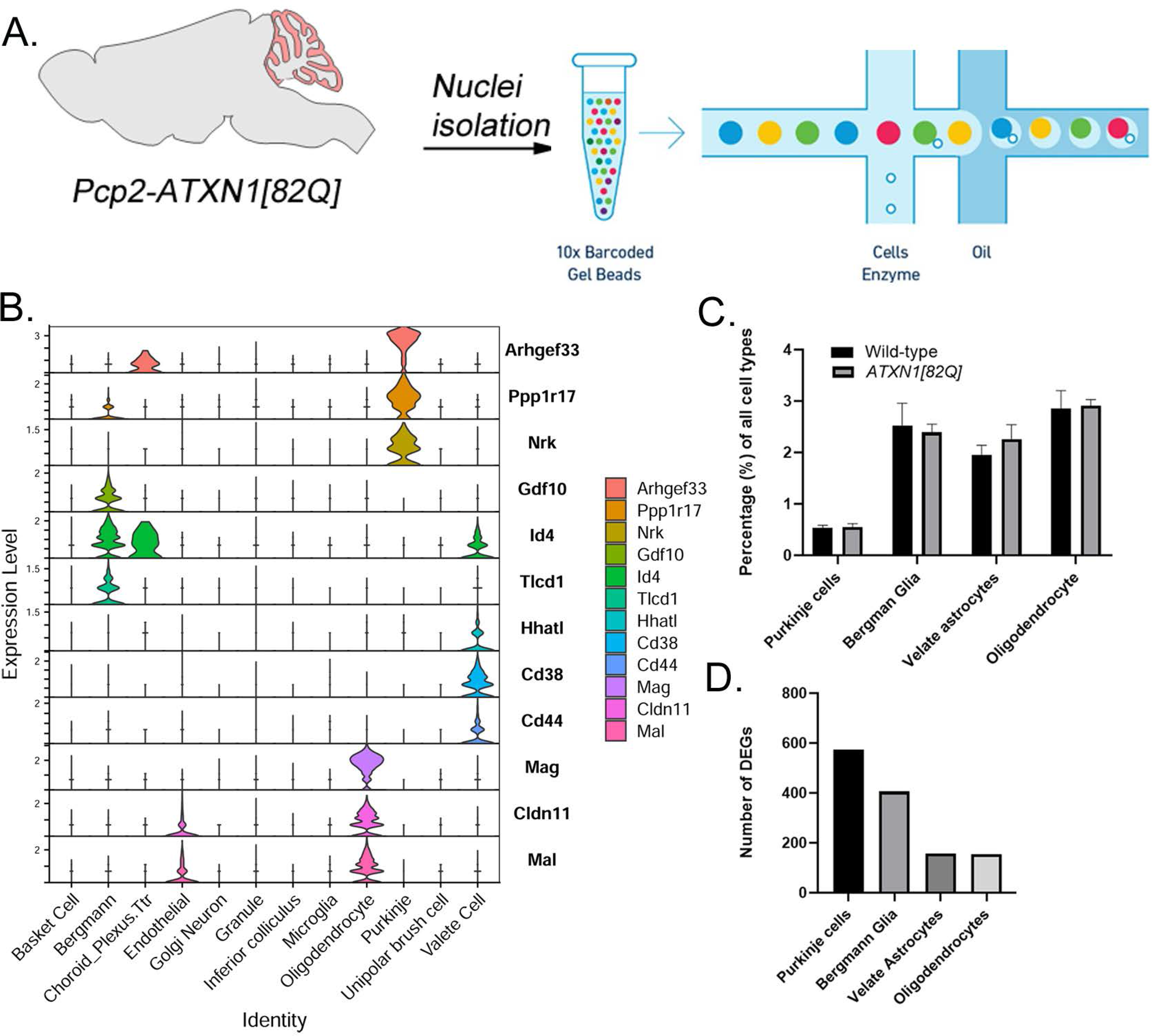
Experimental schematics, cell composition and DEGs per cell type in *Pcp2*-*ATXN1[82Q]* mice. **A.** Nuclei were isolated from the cerebella of 12 weeks old *Pcp2-ATXN1[82Q]* mice and wild-type littermate controls (N=3 of each). After passing library QC nuclei were sequenced on Illumina NovaSeq platform. **B.** Average percentages of Purkinje cells (PC), Bergman Glia (BG), velate astrocytes (VA), and oligodendrocytes in p*cp2-ATXN1[82Q]* mice and wild-type littermate controls (N=3 of each). **C.** Number of differentially expressed genes in each cell type.

We identified a total of 47894 high-quality, snRNAseq profiles. Debris-contaminated nuclei were removed with Seurat using cutoff on number of genes per nuclei or mitochondrial gene percentage, as well as using semi-supervised machine learning classifier Debris Identification using Expectation Maximization (DIEM)[42]. We classified nuclei into the cell types with SingleR (v 1.6.1) using DropViz Cerebellum MetaCells reference [43] (**Figure 1B**).

First, we wanted to determine whether there cellular composition of the cerebellum is altered in SCA1 e.g. whether expression of mutant ATXN1 in Purkinje cells leads to the relative increase or decrease in the numbers of the main cerebellar cell types, including Purkinje cells (PCs), Bergmann glia (BG), velate astrocytes (VA) and oligodendrocytes (OL).

We calculated the percentage of each cell type by dividing the number of specific cell type by the total number of nuclei. As expected majority of the cerebellar nuclei were classified as granule neurons (80.75%). We have not found statistically significant difference in the relative percentages of granule neurons, Purkinje cells, Bergmann glia, velate astrocytes and oligodendrocytes in *Pcp2-ATXN1[82Q]* and wild-type mice, indicating that at this early disease stage we cannot detect a change in the cerebellar cell type composition in *Pcp2-ATXN1[82Q]* mice (**Figure 1C**).

### Expression of mutant ATXN1 in Purkinje cells causes significant gene expression changes in cerebellar glia

Most gene expression studies so far were focused on understanding how mutant ATXN1 affects PCs. Here we extended the analysis of SCA1 gene expression changes to Bergmann glia (BG), velate astrocytes (VA) and oligodendrocytes (OL) in addition to Purkinje cells (PCs) in 12 week old *Pcp2-ATXN1[82Q]* mice [44].

Using differential gene expression analysis we identified differentially expressed genes (DEGs) for each of these four cell types. We found that different cell types have different numbers of DEGS (**Figure 1D**). Remarkably, Bergmann glia and Purkinje cells had a comparable number of DEGs (575 and 406 respectively) at 12 weeks despite the fact that mutant ATXN1 is expressed only in Purkinje cells in these mice. This significant non-cell autonomous transcriptional alterations in Bergmann glia, may suggest that BG are highly sensitive to perturbation in PCs. Both velate astrocytes and oligodendrocytes had a significant number of DEGs (157 and 155 respectively, **Figure 1D**). These results indicate that ATXN1 induced dysfunction in Purkinje cells significantly affects not only nearby Bergmann glia, but also velate astrocytes and oligodendrocytes in a non-cell autonomous manner.

### Molecular profiling of Purkinje cells confirms previously identified molecular pathways and identifies potentially compensatory genes

We compared gene expression changes in PCs to previously reported transcriptional alterations [45][46]. We identified 575 DEGs in Purkinje cells (p.adj q< 0.05), 56% of which were downregulated and 44% upregulated (**Figure 2A**). This prevalence of downregulated genes is consistent with previous results and is generally thought to reflect the cell-autonomous repressive effect of mutant ATXN1 on transcription in PCs [46].

**Figure 2.**
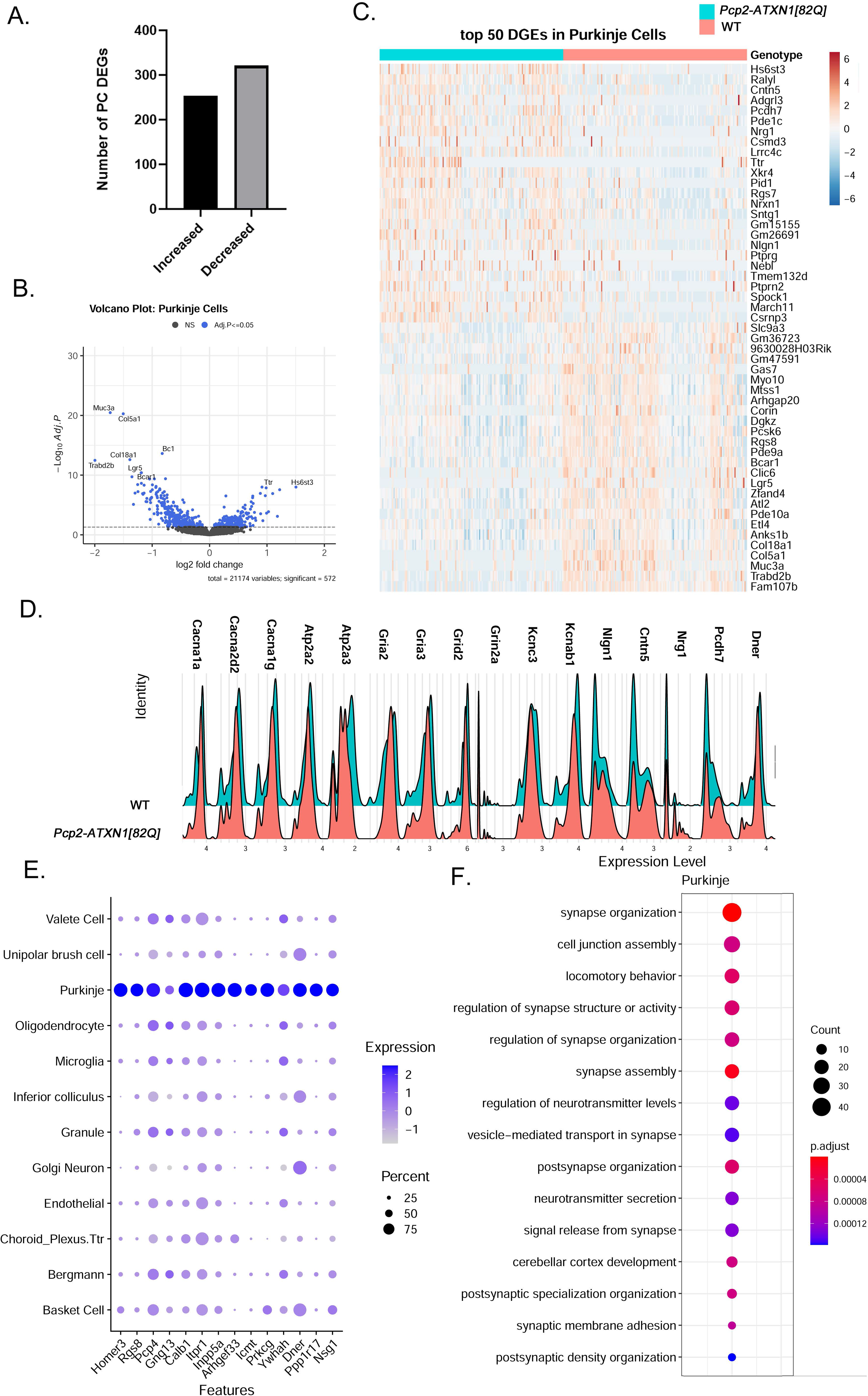
Single nuclei analysis identifies SCA1 disease associated transcriptional changes in Purkinje cells. **A.** Number of upregulated and downregulated DEGs in PCs. **B.** Volcano plot of PCs DEGs **C.** Heatplot of 25 highest upregulated and downregulated PC DEGs. **D.** Ridgeplot showing expression of selected PC DEGs. **E.** Differential expression of Magenta genes in different cerebellar cell types. **F.** Pathway Analysis

Among top ten upregulated genes were *Heparan Sulfate 6-O-Sulfotransferase 3* (*Hs6st3*), *Adhesion G Protein-Coupled Receptor L3* (*Adgrl3*), *Contactin 5* (*Cntn5*), and *Neuregulin 1* (*Nrg1*) known for their roles in neurodevelopment [47][48][49][50]. In addition, SCA1 PCs had increased expression of *RALY RNA Binding Protein-like (Ralyl)* known as Alzheimer’s disease cognitive reserve gene [51] (**Figures 2A-C**). Among upregulated genes were also three vacuolar V-type ATP-ase subunits (*Atp6v0c*, *Atpgv0a1*, and *Atp6v1h*) and *Slc32a1*, encoding vesicular GABA and Glycine Transporter (VGAT) indicating that synaptic vesicles SCA1 PCs may have increased GABA content. These genes may represent compensatory increases in PCs gene expression to restore, or ameliorate synaptic transmission.

Among ten most downregulated genes are four Magenta module genes (*Fam 107b*, *Trabd2b* (regulator of Wnt signaling), *Anks1b*, and *Atl 2)*, three genes encoding extracellular matrix proteins (*Muc3a*, *Col5a1*, *Col 18a*), and two genes encoding proteins involved in GPCR signaling (phosphodiesterase *Pde10* and G protein coupled receptor *Lgr5* that regulates Wnt signaling). Downregulated genes encode calcium voltage gated channels Cav2.1 and Cav3.1 (*Cacna1a, Cacna2d2,* and *Cacna 1g*), calcium endoplasmatic reticulum pumps SERCA1 and SERCA2 (*Atp2a2* and *Atp2a3*), ionotropic glutamate receptors (*Gria2, Gria3, Grid2* associated with Spinocerebellar ataxia autosomal recessive 18, *Grid2ip, Grin2a)*, and potassium channels (*Kcnc3*, associated with SCA13, and *Kcnab1,* associated with Episodic Ataxia Type 1)[52] (**Figures 2C-D**).These results are consistent with previous studies implicating ion channel, calcium and potassium dysregulation and synaptic dysfunction in SCA1 [53].

It is important to note that for most genes, expression in PCs does not seem to be uniform, as indicated by the heatmap and histograms (**Figures 2 C-D**).

Previous studies used weighted Gene Coexpression Network Analysis of the cerebellar bulk RNAsequencing data from *Pcp2-ATXN1[82Q]* mice to identify Magenta Module as a gene network that is significantly correlated with disease progression in Purkinje cells [46]. Thus the Magenta Module is thought to provide a signature of suppressed genes that reflect disease progression in Purkinje cells. We examined which of the Magenta Module genes are changed in Purkinje cells in our single nuclei gene expression analysis. Out of 342 magenta genes Allen brain atlas suggested that 94 (27%) are PC enriched, 175 (51%) PC exclusive and 31 (9%) belonging to multiple cells lines [46]. In previous studies not all of the Magenta Module genes were found to be significantly altered at 12 weeks in *Pcp2-ATXN1[82]* mice. Indeed we found that in our data 103 (30.1%) of Magenta Module genes are altered at 12 weeks of age in *Pcp2-ATXN1[82]* mice. Of those 74 were altered only in PCs (71.8%), while 29 (28.2%) were also changed in two or more cell types, including Bergmann glia, velate astrocytes, and oligodendrocytes. Most intriguing were 13 Magenta Module genes that were changed in all four cell types (**Figure 2E**). For instance expression of gene encoding inositol 3 phosphate receptor 1 (ITPR1) that regulates calcium release from the ER, *Pcp4,* encoding Purkinje Cell Protein 4 that regulate calcium binding to calmodulin, and *Gng13* encoding G Protein Subunit Gamma 13, regulating G protein coupled receptor signaling were significantly suppressed in many cerebellar cell types. This indicates that calcium and G protein signaling are altered not only in PCs, but also in BG, VAs, and OLs. On the other hand expression of *Dner*, encoding for Delta/Notch Like EGF Repeat Containing seems to be altered mostly in cerebellar neurons including PCs, Basket cells, golgi neurons and unipolar brush cells. *Dner* has been previously found to be an important component of PC-BG communication and cerebellar function as expression of DNER in Purkinje cells induces morphological differentiation and functional maturation of Bergmann glia and loss of *Dner* impairs motor behavior of mice [54][55]. It is unclear what role Dner plays in other cerebellar neurons. Pathway analysis of downregulated DEGs using Gene ontology (GO) identified calcium binding (q = 9.8 × 10^−3^), and dendrite and dendritic tree (q values of 1.45 and 1.48 × 10^−5^ respectively), while top Kyoto Encyclopedia of Genes and Genomes (KEGG) pathways of downregulated genes were long-term depression (q = 4.9 × 10^−4^), cholinergic (q = 3.2 × 10^−3^), glutamatergic (q = 3.3 × 10^−3^), serotoninergic (q = 5.2 × 10^−3^) and dopaminergic (q = 5.5 × 10^−3^) synapse and retrograde endocananbinoid signaling (q = 7.1 × 10^−3^) consistent and building up on the previous studies.

Hierarchical enriched pathways analysis of both up and downregulated genes identified many pathways involved in synapse function including synapse organization, cell junction assembly, locomotory behavior, regulation of synapse structure or activity, synapse assembly, and regulation of neurotransmitter levels (**Figure 2F**). Among top GO pathways associated with the upregulated genes were synapse assembly (q = 3.42 × 10^−4^), synaptic membrane adhesion (q = 6.1 × 10^−4^), cell junction assembly (q = 8.8 × 10^−4^), synapse organization (q = 3.2 ×10^−3^) and cell junction organization (q = 8.1 × 10^−3^). Kyoto Encyclopedia of Genes and Genomes (KEGG) identified axon guidance (q = 1.4 ×10^−5^) and synaptic vesicle cycle (q = 2.9 × 10^−3^) as dysregulated pathways.

In conclusion, we confirmed changes in ion transport and synapse dysfunction in SCA1 Purkinje cells as well as identified novel genes that may provide compensatory roles, including *Ralyl*.

### Molecular profiling points to disruptions of multiple pathways in Bergmann glia including Sonic hedgehog signaling from Purkinje cells

Bergmann glia are intimately connected with Purkinje cells with BG cell bodies surrounding the cell bodies of PC and BG radial processes lining up next to PC dendrites and enveloping their synapses in the molecular layer. BG are essential for PCs function and viability through their many roles including maintenance of potassium homeostasis, removal of synaptic glutamate and neurotrophic support [56][57][58][59][60]. Similarly, PCs signaling is important for the development and maintenance of Bergmann glia character [54] [44]. With both their soma and processes next to each other, BG are perfectly poised to sense and respond to PC dysfunction including caused by mutant ATXN1 in SCA1.

We identified 406 DEGs (p.adj <= 0.05) in Bergmann glia. Out of these 151 (37.2%) were downregulated and 255 (62.8%) upregulated (**Figure 3A**). As previous studies indicated that mutant ATXN1 leads to inhibition of gene expression, the increased ratio of upregulated to downregulated genes in BG compared to the opposite ratio in PC, is consistent with the idea of non-cell autonomous gene expression changes in BG.

**Figure 3.**
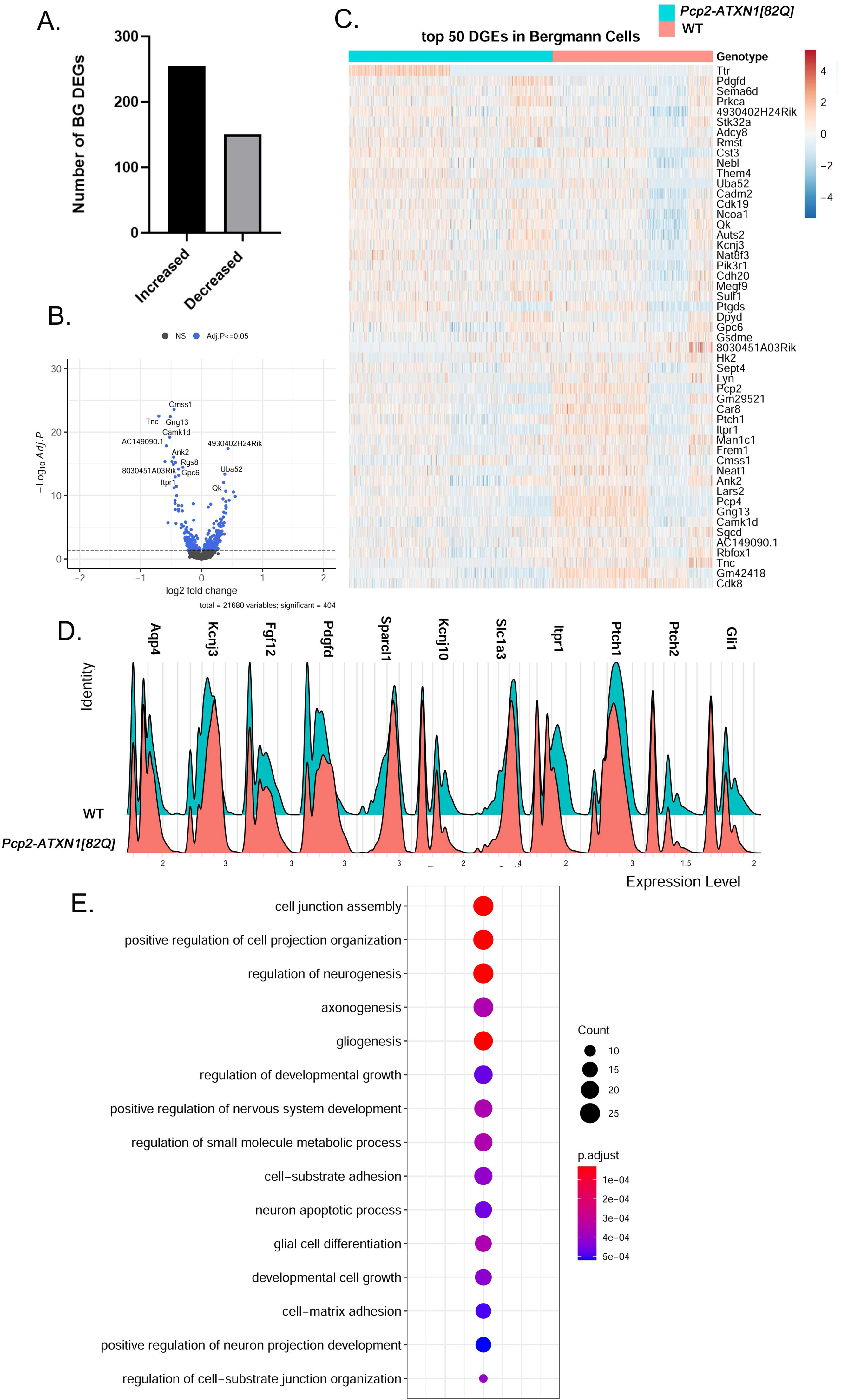
Mutant ATXN1 causes significant non-cell autonomous gene expression perturbations in SCA1 Bergmann glia. **A.** Number of upregulated and downregulated DEGs in BG. **B.** Volcano plot of BG DEGs **C.** Heatplot of 25 highest upregulated and downregulated BG DEGs. **D.** Ridgeplot showing expression of selected BG DEGs. **E.** Pathway Analysis

Among the top upregulated BG genes were transthyretin (*Ttr*) that encodes for the transporter of vitamin A (retinol). Retinol is important for cerebellar function [61] and previous studies found that it is implicated in SCA1 [62]. In addition, BG have increased expression of calmodulin regulated adenylate cyclase (*Adcy8*), that is implicated in Alzheimer’s disease astrocytes and important for memory [63][64], long non coding RNA (lncRNA) *Rmst* that aids in the function of Sox2 transcription factor that is required for BG function [65][66], and *Cystatin* (*Cst3*), the most abundant extracellular inhibitor of cysteine proteases that is thought to be neuroprotective in neurodegenerative diseases[67][68]. We have also found increased expression of *Aquaporin4* (*Aqp4*) that regulates water homeostasis [69] and *Kcnj3*, potassium rectifying channel Kir3.1, that could also be neuroprotective in SCA1. Expression of several trophic factors including fibroblast growth factor (*FGF) 12*, and *Platelet Derived Growth Factor D (PDGFD)* is also increased in BG. Among the upregulated genes was *glial fibrillary acidic protein* (*Gfap*). Additionally, expression of secreted protein acidic and rich in cysteine like 1 (SPARCL1) that directly promotes excitatory synapse formation is increased and is likely compensatory to decreased number of excitatory VGLUT2 synapses on PCs seen at this stage of disease [25]. As increased *Gfap* is thought to reflect during reactive gliosis these BG gene expression changes may implicate increased neuroprotective functions of reactive BG induced by PC dysfunction. (**Figure 3B-C**).

We found a significant changes in the expression of genes involved in homeostatic BG roles that are key for PC neuronal function (**Figure 3D**). Expression of *Kcnj10*, encoding the potassium inward rectifier Kir4.1 is decreased in BG at this stage of disease. Previous studies have found that reduced expression of *Kcnj10* is sufficient to cause PC dysfunction and ataxia in mice [57]. Moreover expression of *Solute carrier family 1 Member 3 (Slc1a3)* that encodes for the glutamate transporter Glast that is key to remove synaptic glutamate is also decreased in BG [70]. These results indicate that SCA1 BG may be less effective in performing these important homeostatic functions.

We also found a large decrease in the expression of inositol tri phosphate receptor 1 (IP3R1), receptor on the ER that upon binding IP3 binding leads to the calcium influx into cytosol [71], that may affect BG function.

Previous studies have found that PC regulate BG character via Sonic hedgehog (Shh) signaling [72]. Expression of Shh receptors *Patched 1 and 2* (*Ptch1, Ptch2*)[73] is decreased in BG in *Pcp2-ATXN1[82Q]* cerebella indicating one critical way in which PC-BG communication is impacted in SCA1 that could affect their function including reduced expression of *Kcnj10* and increased expression of *Aqp4*. Expression of Shh downstream transcriptional activator *Gli1* is also decreased, further supporting dysfunctional Shh PC-BG signaling in SCA1 (**Figure 3D**).

Hierarchical enriched pathways analysis identified pathways involved in development, including cell junction assembly, positive regulation of cell projection organization, regulation of neurogenesis, axonogenesis, gliogenesis, small molecule metabolic processes and neuron apoptotic processes (**Figure 3E**). Among top GO pathways associated with the upregulated genes were nervous system development (q = 1.2 x 10^−9^), regulation of biological quality (q = 1.5 × 10^−9^), cell adhesion (q = 9.8 × 10^−9^), organonitrogen compound metabolic processes (q = 3.8 ×10^−7^) and cell projection organization (q = 5 × 10^−8^). KEGG identified circadian entrainment (q = 3.6 × 10^−2^) as dysregulated pathway in BG. This is very intriguing considering the role of cerebellum in sleep and sleep disturbances in SCA patients[74][75]. Go pathway analysis of downregulated genes identified transmembrane transporter binding (q = 7 × 10^−3^), smoothened binding and hedgehog receptor activity (q = 4.9 × 10^−2^), cell cell signaling (q = 6.4 × 10^−7^), apoptotic processes (q = 1.3 ×10^−2^) and synapse (q = 5 × 10^−14^). KEGG analysis of downregulated genes identified retrograde endocannabinoid signaling (q = 1 × 10^−3^), glutamatergic (q = 2 × 10^−3^) and dopaminergic synapses (q = 4.3 × 10^−2^), long-term depression (q = 8.1 × 10^−3^) and spinocerebellar ataxias (q = 8.4 × 10^−3^). Thus pathway analysis further supports that SCA1 BG increase their neuroprotective functions to compensate for PC dysfunction, but they decrease their homeostatic roles that may promote PC dysfunction. This decrease in the expression of homeostatic genes may be in part due to reduced ability to receive Shh signaling from PC due to the reduced expression of *Ptch1* and *Ptch2*.

### Increased expression of genes to promote neurogenesis, gliogenesis and synaptogenesis indicate beneficial compensatory gene expression changes in velate astrocytes

Velate astrocytes are type of cerebellar astrocytes that reside in the granule cell layer. As such they are surrounded by the most numerous neurons in the brain and are the only astrocytes in the brain that are largely outnumbered by neurons [76]. Moreover while volume of the whole cerebellum is reduced in SCA1, much less is known about the pathology in granule cell layer.

We have identified 157 DEGs (p.adj <= 0.05) in SCA1 velate astrocytes compared to their wild-type littermate controls with 114 or 72.6% of genes upregulated (**Figure 4A, D**). One of genes with increased expression was *Vimentin* (*Vim*) encoding for the intermediate filament predominantly found in astrocytes and upregulated when they are undergoing reactive astrogliosis [77][78]. This result may indicate that VA undergo reactive gliosis in SCA1 (Figure 4D). Expression of *apolipoprotein E* (*ApoE*) was also significantly increased in SCA1 velate astrocytes. As ApoE, among other roles contributes to astrocyte activation and BDNF secretion, this increase in ApoE may contribute to VA activation and increased BDNF early in SCA1. [79][80]. Among the top upregulated genes was *Transthyretin* (*Ttr*), encoding for protein primarily responsible for the transport of thyroxin and retinol-retinol binding complex (RBP-complex) but also plays a role in oxidative stress [81]. Moreover increased expression of *Ttr* is suggested to play a role in resistance to AD neurodegeneration [82]. *Neural Cell adhesion molecule 2* (*Ncam2*) encodes NCAM2 protein that is involved in neuronal migration, cell positioning during corticogenesis, neuronal differentiation and synaptogenesis during development and in the calcium signaling, maintenance of presynaptic and postsynaptic compartments in adult brains. In addition, it has been proposed that a decrease in NCAM2 levels is associated with loss of structure of synapses in the early stages of neurodegenerative diseases [83]. Recently, *CUB And Sushi Multiple Domains 1*(*Csmd1*) was suggested to oppose the complement cascade that facilitates synaptic loss in neurodegeneration[84]. Therefore, it is reasonable to assume that increased expression of *ApoE*, *Ttr*, *Ncam2*, and *Csmd1* may be neuroprotective in SCA1 as well.

**Figure 4.**
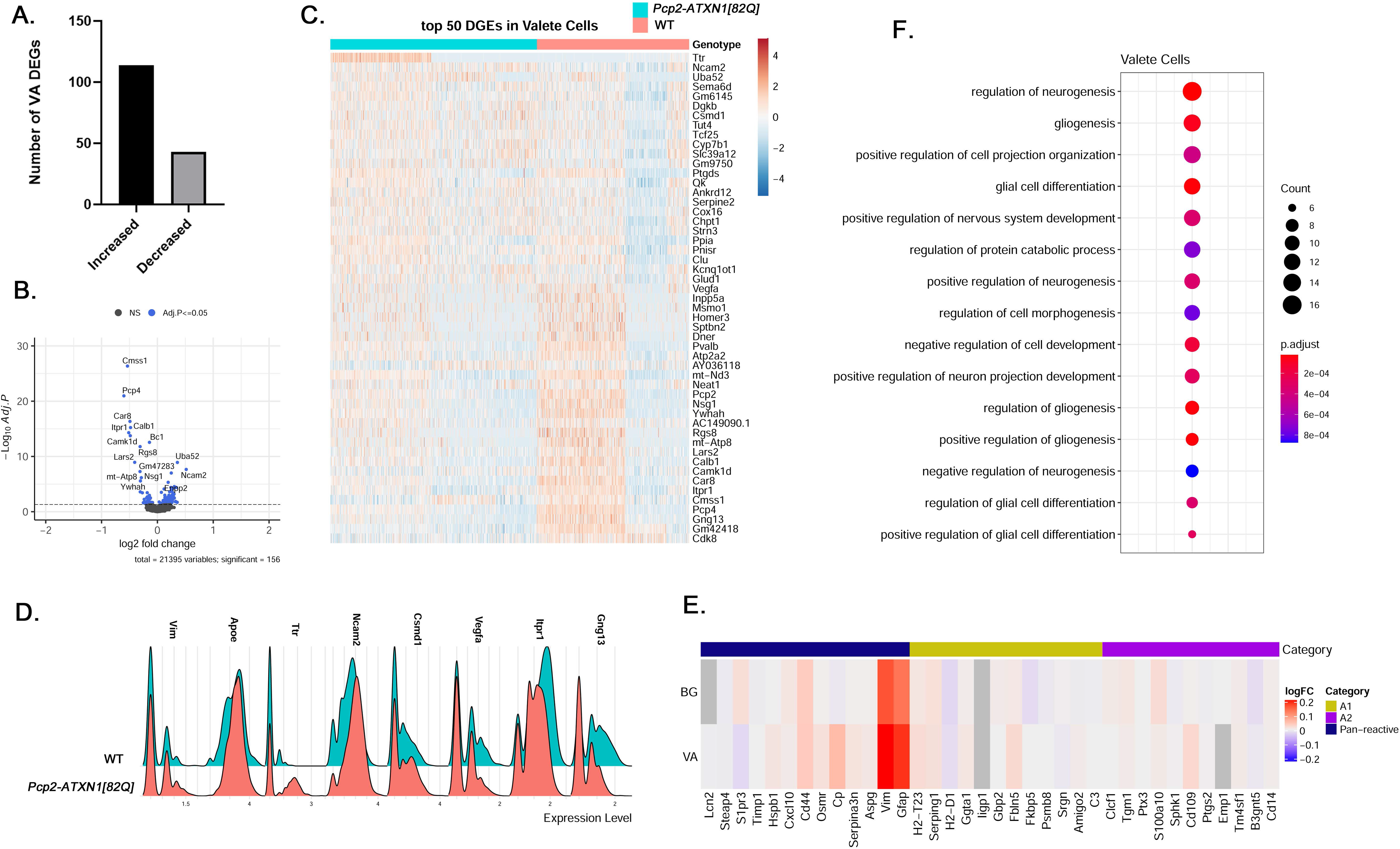
Velate astrocytes gene expression changes in SCA1. **A.** Number of upregulated and downregulated DEGs in VA. **B.** Volcano plot of VA DEGs **C.** Heatplot of 25 highest upregulated and downregulated VA DEGs. **D.** Ridgeplot showing expression of selected VA DEGs. **E.** Heatplot showing expression of A1/A2 genes in BG and VA. **F.** Pathway Analysis

Most of the homeostatic astrocytic genes were not altered with the exception of reduced expression of *vascular endothelial growt*h *factor* (*VEGF)*, neurotrophic factor whose decrease was previously shown to contribute to Purkinje cell pathology in SCA1 [85] and genes encoding astrocyte receptors *Itpr1* and *G Protein Subunit Gamma 13* (*Gng13*), that regulate calcium and GPCR signaling were downregulated (**Figure 4D**).

Finally, we examined the expression of A1 and A2 astrocyte genes in VA and BG. Previous study identified two different types of reactive astrocytes that were termed “A1” and “A2” respectively. A1 astrocytes are thought to be harmful as they up-regulate classical complement cascade genes previously shown to be destructive to synapses, while A2 astrocytes are thought to be neuroprotective due to increased expression of neurotrophic factors. We found increased expression of panreactive genes Gfap, Vim, and CD44 but have not found any clear pattern or distinction in the expression of A1 or A2 genes (**Figure 4E**)

Pathways analysis identified developmental pathways including regulation of neurogenesis, gliogenesis, positive regulation of projection organization, regulation of nervous system development as well as, regulation of protein catabolic process (**Figure 4F**).

Pathway analysis of the upregulated identified many developmental pathways including cell morphogenesis, regulation of neuron projection development, positive regulation of gliogenesis, and glial differentiation (q values of 3.2 X10^−6^, 1.5 X10^−5^, 3.9 X10^−5^, and 4.5 X10^−5^ respectively).

Pathway analysis of the downregulated genes identified calcium ion binding (q = 3.1 X10^−3^), vesicle mediated transport in synapse (q = 3.5 X10^−2^), negative regulation of neuronal death (q = 4.1 × 10^−2^), glutamatergic synapse (q = 4.5 ×10^−4^), spinocerebellar ataxia (q = 1.3 ×10^−3^), retrograde endocannabinoid signaling (q = 2.4 ×10^−2^), and long-term depression (q = 2.7 ×10^−2^).

### Gene expression analysis indicates reactive activation and perturbation in calcium homeostasis in SCA1 oligodendrocytes while pathways analysis revealed association with locomotor behavior and cognition

Oligodendrocytes (OL) envelop axons with myelin and maintain long-term axonal integrity [86]. Previous studies demonstrated atrophy in white matter already present in pre-manifest carriers of SCA1 mutation [29] as well as in *Pcp2-ATXN1[82Q]* mice [30]. These studies indicated functional changes in oligodendrocytes in SCA1 from the early stages of disease progression. Moreover recent sn-RNAseq study indicated decreased expression of mature oligodendrocyte genes, including *Myelin Oligodendrocyte Glycoprotein* (*Mog*) and oligodendrocyte precursor (OPC) genes involved in OL formation in a knock-in mouse model of SCA1, *Atxn1^154Q/2Q^* mice. In *Atxn1^154Q/2Q^* mice mutant Atxn1 is expressed widely. To gain insight into which of these genes are induced in response to PC pathogenesis, we examined genes and pathways altered in oligodendrocyte nuclei from *Pcp2-ATXN1[82Q]* mice compared to littermate controls.

We found 154 DEGs in oligodendrocytes in *Pcp2*-*ATXN1[82Q]* cerebella, with majority (94 DEG or 61%) of genes being upregulated (**Figure 5A**).

**Figure 5.**
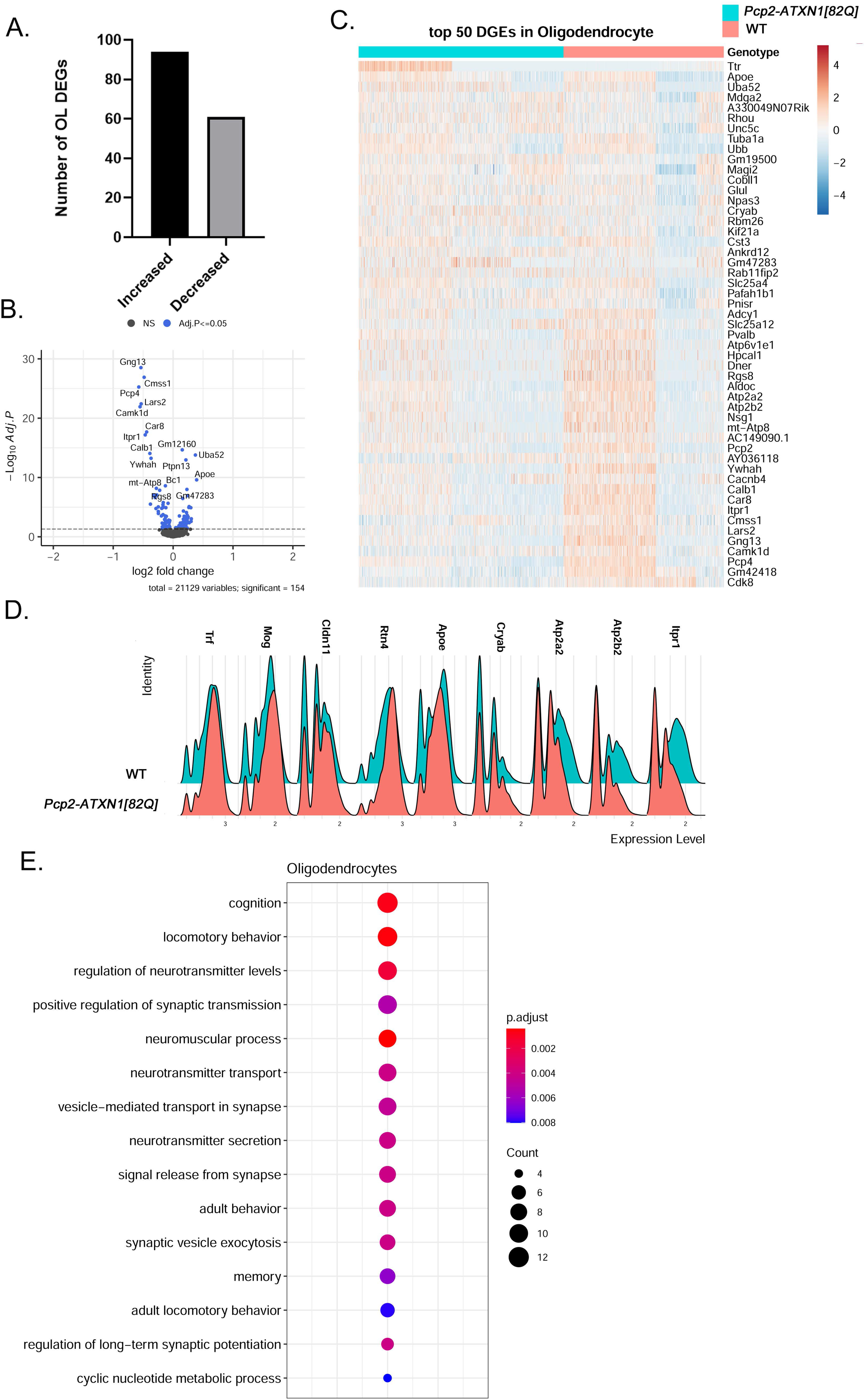
Gene expression changes in SCA1 oligodendrocytes. **A.** Number of upregulated and downregulated DEGs in OL. **B.** Volcano plot of OL DEGs **C.** Heatplot of 25 highest upregulated and downregulated OL DEGs. **D.** Ridgeplot showing expression of selected OL DEGs. **E.** Pathway Analysis

While OL numbers are not altered, expression of several oligodendrocyte marker genes including *Transferrin* (*Trf)*, *Mog*, *Claudin 11* (*Cldn11*) and *Reticulon 4* (*Rtn4*) was increased (**Figures 5B-D**)[86].

We found altered expression of many genes important for oligodendrocyte function (Figure 5E). Among them is increased expression of *Glutamate-Ammonia Ligase* (*Glul*) encoding for glutamine synthetase (GS) a key enzyme that catalyzes the conversion of glutamate to glutamine. Previous studies demonstrated that OL glutamine synthetase is important for glutamatergic transmission [87], and that *Glul* expression in OL is increased in chronic pathological conditions including amyotrophic lateral sclerosis (ALS) and multiple sclerosis (MS)[88]. We also found increased expression of *Crystallin Alpha B* (*Cryab*) and *ApoE* in SCA1 OL. Increased *Cryab* and *ApoE* have been associated with protective effects and astrocyte gliosis in other neurodegenerative diseases including Parkinson’s disease (PD), Alzheimer’s disease (AD) and MS [89][90][91][92][80]. Increased expression of transcriptional factor *JunD*, a functional component of activated protein 1 (AP1) complex that regulates many of the inflammatory and immune genes, also indicates reactive SCA1 OL response in pathogenesis [93].

Among downregulated genes were *Atp2a2* and *Atp2a3* encoding for calcium endoplasmatic reticulum pumps SERCA1 and SERCA2, *Atp2b2* encoding for Plasma Membrane Calcium-Transporting ATPase, and *Itpr1* implicating perturbation in calcium homeostasis in reactive SCA1 OLs (**Figures 5 B-D**).

Intriguingly top two pathways identified were cognition and locomotor behavior, implicating that molecular pathway alterations in SCA1 oligodendrocytes are associated with the changes in cognition and locomotion in these mice. Pathway analysis also identified regulation of neurotransmitter levels, positive regulation of synaptic transmission, vesicle-mediated transport in synapses and neurotransmitter secretion potentially implicating OL in alterations of synaptic transmission in SCA1 (**Figure 5E**).

Pathway analysis of downregulated genes identified pathways involved in transport and binding of ions including inorganic cation transmembrane transporter activity (q = 7.5 X10^−6^), calcium binding (q = 9.3 X10^−6^), P-type calcium transporter activity (q = 4.2 X10^−4^), synaptic signaling including synaptic signaling (q = 1.6 × 10^−7^), modulation of chemical synaptic transmission (q = 5.4 × 10^−6^), glutamatergic synapse (q = 6.4 ×10^−6^), and Spinocerebellar ataxia (q = 2.9 ×10^−5^). Among the pathways identified with upregulated genes were movement of cell or subcellular component (q = 9.4 X10^−5^), cell migration (q = 8.8 X10^−4^), myelin sheath (q = 2.4 X10^−7^), synapse (q = 1.4 X10^−4^), Parkinson’s disease (q = 4.6 X10^−3^), and iron uptake and transport (q = 3.8 X10^−2^).

## Discussion

Here, we have employed single nuclei RNA sequencing to investigate transcriptional changes in individual cerebellar cell types during the early stages of disease progression in transgenic Purkinje cell specific mouse model of SCA1 *Pcp2-ATXN1 [82Q]* line. These results provide insight into how pathogenic processes within neurons, in this case Purkinje cells, affect gene expression and signaling pathways within cerebellar astrocytes and oligodendrocytes increasing our understanding of the causal sequences of progressive dysfunctions during early stages of disease.

There are several important findings from our study. First, mutant ATXN1 expression in Purkinje cells causes profound transcriptional alterations in cerebellar glia in a non-cell autonomous manner. Majority of the altered glial genes were upregulated (Bergmann glia (62.8%), velate astrocytes (72.6%) and oligodendrocytes (61%)). On the other hand mutant ATXN1 primarily caused downregulation of gene expression in cell autonomous manner in PCs as 56% of genes had reduced expression [94]. At 12 weeks there is no Purkinje cell loss in *Pcp2-ATXN1[82Q]* so we propose that these glial transcriptome changes are in response to Purkinje cell dysfunction [46]. In addition, we found that proximity to dysfunctional neurons seems to be an important factor in determining the extent of the glial non-cell autonomous transcriptional perturbations in SCA1. Bergmann glia that reside next to Purkinje cells have similar number of DEGs as Purkinje cells themselves and approximately three times more DEGs than velate astrocytes and oligodendrocytes that reside further away from PCs. This finding supports the previous evidence of very intimate relationship between PCs and BG [95].

Second, we identified transcriptional changes in PCs, BG, VA and OL that may be relevant to disease pathogenesis. For PCs we validated the findings from previous studies but also identified novel, potentially compensatory gene expression changes such as increased expression of *Ralyl*, *Atp6v0c*, *Atpgv0a1*, *Atp6v1h*, Atp6v and *Slc32a1. Ralyl* is an RNA binding protein known for its neuroprotective role in Alzheimer’s [51]. Increased expression of three vacuolar V-type ATP-ase subunits (*Atp6v0c*, *Atpgv0a1*, and *Atp6v1h*) that provide electrochemical gradient to fill synaptic vesicles with GABA and *Slc32a1*, encoding vesicular GABA and Glycine Transporter (VGAT) that transports GABA in the vesicles may suggest increased GABA content in PCs synaptic vesicle. As spontaneous firing rate of PCs is decreased in *Pcp2-ATXN1[82Q]* mice [96], this increase in GABA per vesicle may represent compensatory change to restore, or ameliorate synaptic transmission by increasing the amount of GABA released at PC terminals.

We validated previously reported decrease in the expression of glutamate transporter *Slc1a3* in Bergmann glia [70], and identified additional perturbations in the expression of homeostatic genes such as *Kcnj10* that are critical for PC function [97][57]. Moreover, our results suggest perturbed Shh signaling as one possible mechanism of these BG molecular alterations. Sonic hedgehog (Shh) signaling is one of the key communication pathways between PCs and BG involved in regulation of BGs character by PCs [72]. We found that the expression of Shh receptors Patched 1and 2 is decreased in BGs. Moreover, the expression of Shh signaling downstream transcription factor Gli1 is also reduced on SCA1 BG. Previous studies have shown that SHH-Ptch2 communication regulates expression of homeostatic genes *Slc1a3*, *Kcnj10*, and *Aqp4* in Bergmann glia [44]. Therefore it is possible that decreased Shh signaling contributes to reduced expression of S*lc1a3* and *Kcnj10* and increased expression of *Aqp4* that we found in SCA1 BG. Thus reduced PC-BG communication via Shh signaling may impair expression of BG identity genes and their ability to support Purkinje cells by maintaining homeostasis of potassium, glutamate and water, leading to progressive cerebellar dysfunctions.

In velate astrocytes we identified increased expression of several genes such as *ApoE, Ncam2*, and *Csmd1* that may provide neuroprotection. For instance we have previously demonstrated that BDNF provides neuroprotection in SCA1 [98][99]. As ApoE, among other roles contributes to astrocyte activation and BDNF secretion, this increase in ApoE may contribute to VA activation and increased BDNF that we have seen early in SCA1 [79][80]. *Ncam2* and *Csmd1* are protective against loss of synapses [83][84], and increase in their expression in SCA1 VA may ameliorate loss of synapses seen in SCA1.

Perhaps most intriguing are the transcriptional changes we identified in oligodendrocytes. While OL numbers do not seem to be altered at 12 weeks, we have found increased expression of OL marker genes and genes indicating OL reactive activation including *Trf*, *Mog*, *Cldn11*, *Rtn4*, *JunD, Cryab* and *ApoE*.

Third, we identified shared DEGs and pathways that are altered in all four cell types in SCA1. These shared pathways are involved in regulation of synapses, cytoarchitecture, calcium and development. For instance several magenta genes and Ttr are altered in PCs, BG, VA and OL. Transthyretin (Ttr) is a protein that binds and distributes thyroid hormones and retinol. Previous studies identified the importance of retinol in PC pathogenesis in SCA1 [62]. Moreover, recent studies found that it plays in regulation of OPCs proliferation and survival [100], described it as regulator of the δ-GABAA-R expression in cerebellar granule neurons [101] and regulator of glycolysis in astrocytes [102]. Therefore increased Ttr expression could be beneficial for many cerebellar cell types in SCA1. It will be important to understand the mechanism of *Ttr* upregulation in different cerebellar cell types and which cells are affected by increased *Ttr* in SCA1 cerebella. In addition, several Magenta genes, including genes involved in calcium homeostasis, such as *Atp2a2* and *Itpr1* are reduced in all four cell types, indicating that calcium signaling may be altered in SCA1 PCs, BG, VA and OL. Identification of shared DEGs and pathways indicate that in response to the neuronal dysfunction neurons and glial cells undergo similar perturbations early in disease progression, prior to manifestation of motor symptoms. Thus these shared pathways may be ideal targets for therapeutic approaches.

It is likely that some of the molecular changes we identified are compensatory allowing for continued cerebellar function, and that some may be pathogenic, promoting disease progression. Our results provide a framework to investigate how individual pathogenic processes contribute to the sequence of progressive cerebellar dysfunctions in SCA1. Our results are freely available and we hope that they will stimulate future studies to causally investigate how these gene and pathway perturbations contribute to SCA1 pathogenesis as well as facilitate development of novel therapeutic approaches.

## Acknowledgements

We acknowledge all the members of Orr and Cvetanovic laboratories for thoughtful discussions and feedback on the study. Work in this study was aided by the Single Cell Services of the University of Minnesota Genomics Center.

## Author Contributions

M.C. conceptualized the study. E.B. performed experiments. Y.Z. and M.C. analyzed data. M.C., E.B. and YZ wrote the manuscript.

## Figure legends

**Supplementary Figure 1. Pathway analysis of Magenta Module genes altered in PCs, BG, VA and OLs in *Pcp2-ATXN1[82Q]* mice.**

