## Supplementary figures and images for "Single-nuclei RNA sequencing uncovers non-cell autonomous changes in cerebellar astrocytes and oligodendrocytes that may contribute to Spinocerebellar Ataxia Type 1 (SCA1) pathogenesis"

### Supplementary Figure 1

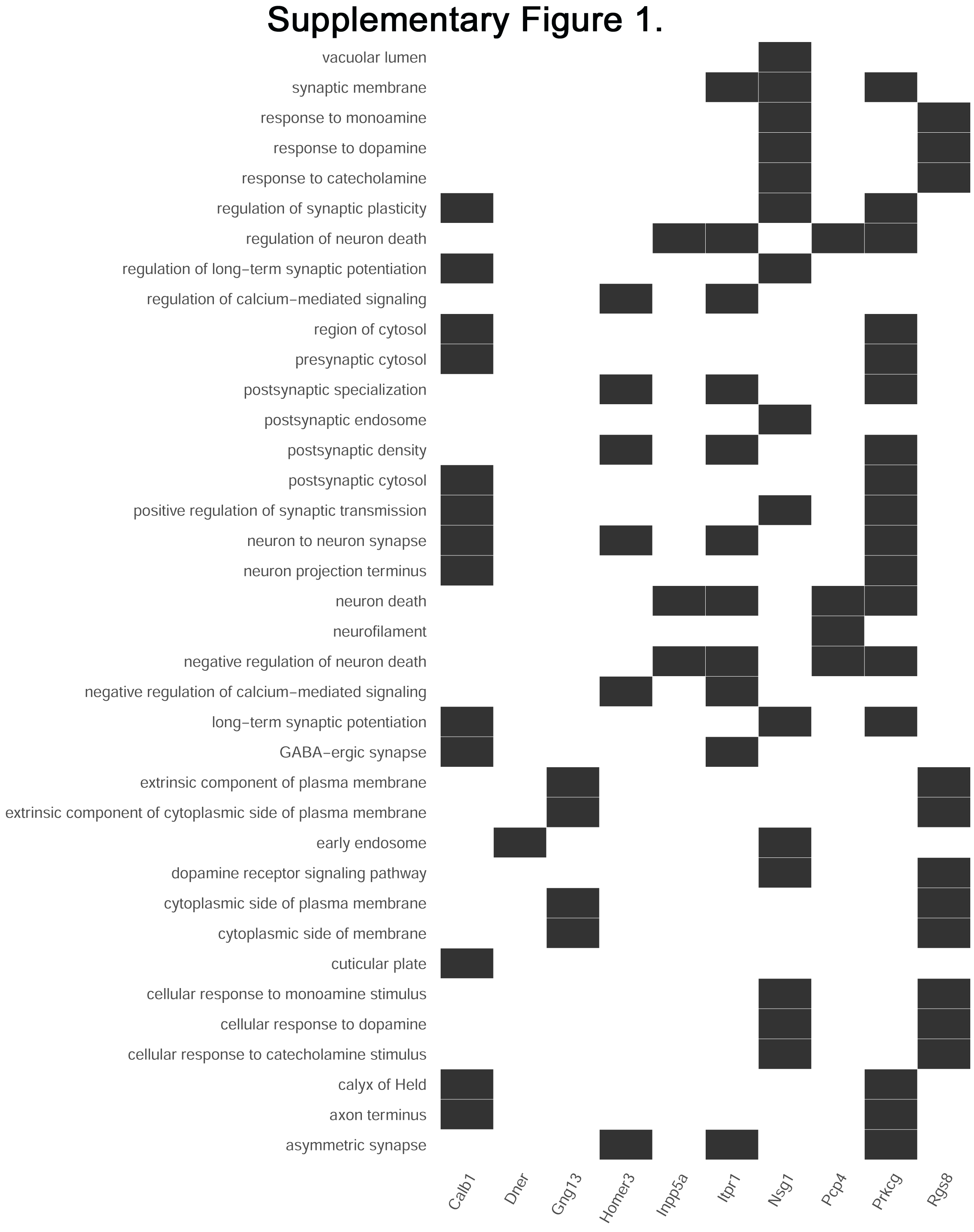
